# Integrated epidemiological and genomic analysis of some respiratory Bovine Coronavirus isolates reveals circulation of GIIb strains and ongoing viral evolution in U.S. Cattle (2020–2025)

**DOI:** 10.64898/2026.03.12.711484

**Authors:** Abid Ullah Shah, Csaba Varga, Philip Guger, Maged Gomaa Hemida

**Author notes:** Equal contribution (Abid Ullah Shah, Csaba Varga). **Corresponding author: Maged Gomaa Hemida,** Department of Veterinary Biomedical Sciences, Lewyt College of Veterinary Medicine, Long Island University, Brookville, NY, 11548.

## Abstract

Bovine coronavirus (BCoV) is an important contributor to the respiratory disease complex in cattle; however, integrated genomic and epidemiological data describing currently circulating respiratory BCoV strains in the United States remain limited. The objective of this study was to monitor respiratory BCoV at the genomic level and analyze its epidemiological patterns over a five-year period. A total of 4,505 respiratory samples submitted to a diagnostic laboratory between January 2020 and November 2025 were analyzed, of which 693 (15.38%) tested positive for BCoV. Positivity was highest in young calves (0–40 days; 20.0%) and declined significantly with increasing age based on logistic regression analysis. Temporal trend analysis using LOESS smoothing and the Mann–Kendall test showed no significant monotonic change in BCoV detection during the study period. Co-infection analysis indicated that BCoV was commonly detected with other viral respiratory pathogens, while bacterial pathogens predominated in many samples. Lung tissues from infected cattle were screened by RT-PCR, and selected samples with high viral loads were subjected to next-generation sequencing. Complete genome sequencing identified four respiratory BCoV isolates (∼31 kb), all clustering within genotype GIIb with recent U.S. strains. Comparative genomic analysis revealed several amino acid substitutions in structural and non-structural proteins that may influence viral attachment, replication, and tissue tropism. These findings provide updated epidemiological and genomic insights into respiratory BCoV circulating in U.S. cattle.

## Introduction

BCoV is an endemic Betacoronavirus of cattle in many regions across the globe with major implications for animal health, livestock productivity, and coronavirus evolution (1–3). BCoV is a well-established etiological agent of neonatal calf diarrhea, winter dysentery in adult cattle, and respiratory disease across age groups, where it functions as an important viral component of the bovine respiratory disease complex (BRDC) (4–6). Despite its long-standing recognition, BCoV continues to circulate extensively in cattle populations worldwide and remains under continuous evolutionary pressure, resulting in genetic diversification that may influence tissue tropism, pathogenicity, and transmission dynamics (7–11).

BCoV belongs to the species Betacoronavirus 1, which also includes human coronavirus OC43 (HCoV-OC43), and other coronaviruses affecting various species of animals (12–14). Phylogenetic and molecular clock analyses strongly support a historical bovine-to-human spillover event that gave rise to HCoV-OC43, underscoring cattle as a critical reservoir of Betacoronavirus diversity and highlighting the broader One Health relevance of BCoV surveillance (15–19). Nevertheless, contemporary molecular data on BCoV particularly respiratory isolates remain limited, constraining our understanding of how ongoing viral evolution intersects with disease expression in cattle (10, 20, 21).

From an epidemiological perspective, respiratory BCoV is frequently detected in calves and young cattle presenting with respiratory disease and is commonly identified alongside viral and bacterial co-pathogens associated with BRDC (22–24) . Age-dependent susceptibility has been repeatedly observed, yet most studies rely on categorical age groupings and short surveillance windows, limiting quantitative assessment of age as a continuous risk factor (3, 25, 26). Longitudinal analyses that evaluate temporal trends in BCoV detection over multiple years are also scarce, particularly those extending into the post-2020 era of intensified molecular diagnostics (27, 28).

At the molecular level, BCoV possesses a ∼31 kb positive-sense RNA genome encoding the replicase polyprotein (ORF1ab), four major structural proteins spike (S), hemagglutinin-esterase (HE), membrane (M), and nucleocapsid (N) and several accessory non-structural proteins. The spike glycoprotein is the primary determinant of viral attachment, receptor engagement, and tissue tropism, whereas the HE protein mediates reversible binding to 9-O-acetylated sialic acids and facilitates viral release. Accumulating evidence indicates that amino acid substitutions within the S1 domain of spike and within functional regions of HE can alter viral infectivity, antigenicity, and potentially respiratory versus enteric tropism. However, most available genomic data have been generated from enteric samples, leaving respiratory BCoV isolates, especially those derived directly from lung tissue, substantially underrepresented.

In addition to the major structural proteins, the accessory non-structural proteins encoded between the spike and membrane genes, including the 4.8 kDa, 4.9 kDa, and 12.7 kDa proteins, represent some of the least understood components of the BCoV genome. Truncations, insertions, and the emergence of novel open reading frames in this region have been sporadically reported and may reflect host adaptation, genome plasticity, or functional redundancy. Comprehensive whole-genome sequencing is therefore essential to capture the full spectrum of genetic variation across both conserved and rapidly evolving regions of the BCoV genome (29–31).

The advancement of the next generation sequencing (NGS) technology enriched our knowledge about the genome structures and organization of various viral diseases affecting various species of animals and birds including cattle (7, 32–36).

Importantly, integrating molecular characterization with large-scale epidemiological analysis provides a powerful framework for understanding how viral evolution manifests at the population level. Advances in diagnostic testing and sequencing capacity since 2020 offer a unique opportunity to examine respiratory BCoV circulation across multiple years, production systems, and age groups while simultaneously resolving genomic changes that may influence viral fitness and disease expression. Such integrative approaches remain uncommon for endemic livestock coronaviruses (37–39).

In this study, we provide a comprehensive epidemiological and molecular portrait of contemporary respiratory BCoV circulation in the United States (U.S.). Our findings reveal sustained circulation of genotype GIIb viruses and identify multiple novel amino acid substitutions across structural and non-structural proteins, underscoring ongoing viral evolution in respiratory tissues. Together, these results advance understanding of respiratory BCoV biology and highlight the importance of integrated genomic surveillance to inform diagnostics, vaccine development, and disease control strategies.

## 2. Materials and Methods

### 2.1. Study data and variables

The study included reverse transcription real-time PCR (RT-PCR) test results for bovine coronavirus (BCoV) obtained from specimens representing the respiratory tract of cattle submitted to the Iowa State University Veterinary Diagnostic Laboratory (ISU VDL) between January 2020 and November 2025. Diagnostic submissions to the ISU VDL and associated metadata were selected from the database if the submission represented only bovine species of any age, included lung tissues evaluated for histopathological lesions, reported RT-PCR results from the ISU VDL bovine respiratory panel (BRP: targets bovine coronavirus, bovine viral diarrhea virus1/2, bovine respiratory syncytial virus, and bovine herpesvirus type 1, *Mannheimia haemolytica*, *Histophilus somni*, *Mycoplasma bovis*, and *Pasteurella multocida*), and was diagnosed with respiratory disease involving at least one infectious etiology. Diagnostic submissions satisfying these criteria may have additional metadata, including bacterial culture results on lung tissue and immunohistochemistry (IHC). Data were reported in an Excel spreadsheet for descriptive statistics. Additional statistical analyses were conducted using R software (Version 4.1.2; R Foundation for Statistical Computing, Vienna, Austria) within the RStudio platform.

### 2.2. Descriptive Statistics

The proportion of positive test results was calculated overall by dividing the total number of positive samples by the total number of samples tested. In addition, the proportion of positives by age groups for 4 age categories (0–40 days, 41–140 days, 141–240 days, 241+ days) by location (State) of submissions, and by production type (cow/calf, feedlot, dairy, and others) was also calculated.

### 2.3. Trend analysis

A trend analysis was conducted to assess the proportion of positive samples over the study period (January 2020 to November 2025). A locally estimated scatterplot smoothing (LOESS) method was applied to smooth the data. A smoothing line was added to the image to visualize the underlying trend and assess the trend in the proportion of positive results over time. This non-parametric method is useful for visualizing trends in data that may be nonlinear. To assess whether there was a statistically significant monotonic trend in the data, the Mann-Kendall (MK) test was applied. This non-parametric test evaluates the null hypothesis of no monotonic trend against the alternative hypothesis of a monotonic trend. The result of the MK test is reported as a tau value and a 2-sided p-value. The positive values indicate an increase in sample positivity over time, while a negative value indicates a decrease.

### 2.4. Assessing co-infections

The following pathogens were included in the analysis: Bovine coronavirus (BCoV), Bovine herpesvirus 1 (also known as infectious bovine rhinotracheitis virus) (BHV-1), Bovine respiratory syncytial virus (BRSV), Bovine viral diarrhea virus (BVDV); bacterial pathogens: *Histophilus somni* (formerly *Haemophilus somnus*) (*H. somnus*), *Mycoplasma bovis* (*M. bovis*), *Mannheimia haemolytica* (*Mannheimia*), *Pasteurella multocida* (*Pasteurella*). Only samples with complete test results across all pathogens were included. Test outcomes were standardized to positive or negative, and indeterminate or missing results were excluded. For downstream analyses, pathogen detection status was converted to a binary format, with positive detections coded as 1 and negative detections coded as 0, generating a binary co-infection matrix for all samples.

To quantify similarity among samples based on their co-infection profiles, pairwise dissimilarities were calculated using Gower distance. Pathogen variables were specified as symmetric binary variables, allowing both shared presences and shared absences to contribute to similarity estimates. Hierarchical agglomerative clustering was subsequently performed on the resulting Gower distance matrix using the average linkage method, and a dendrogram was constructed to display the overall structure of co-infection patterns among samples.

To identify the optimal number of clusters, silhouette analysis was conducted on cluster solutions ranging from two to ten. For each candidate number of clusters (k), samples were assigned to clusters using dendrogram cutting, and silhouette widths were calculated based on the Gower distance matrix. The average silhouette width, which reflects both within-cluster cohesion and between-cluster separation, was computed for each k. The optimal number of clusters was defined as the value of k that maximized the average silhouette width. For each identified cluster, the number and proportion of samples were calculated. Additionally, cluster-specific summaries were generated to quantify the number and proportion of samples testing positive for each pathogen within a cluster.

### 2.5. Logistic Regression Analysis

A logistic regression model was fitted to examine the relationship between age and the probability of testing positive for bovine coronavirus. The model was specified as:

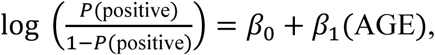

where *P(positive)* is the probability of a positive test result, and age is the continuous predictor. The model coefficient was estimated using maximum likelihood estimation, assuming a binomial distribution with a logit link function, and the odds ratios (ORs) were calculated by exponentiating the regression coefficients. Results were reported as odds ratios with corresponding 95% confidence intervals. Statistical significance was assessed using Wald tests, with a significance level of α = 0.05. Model diagnostics were conducted to assess the fit and adequacy of the logistic regression model. Predicted probabilities were generated for the entire range of age to examine the relationship between age and the probability of a positive result.

### 2.6. Sample collection and processing

A total of thirteen tissue suspensions from bovine lungs were received from the ISU-VDL. Samples were collected from BCoV RT-PCR bovine lungs between 2023 and 2024. All samples were tested for bovine viral infections, including BVDV1, BVDV2, BRSV, and IBRV, using RT-PCR. The samples positive for any of these viruses were excluded from further analysis. The tissue suspension was processed as described previously (31) and stored at -80 °C for further analysis.

### 2.7. Viral RNA extraction and BCoV confirmation by RT-PCR

Total RNA from tissue suspension was isolated using TRIzol LS reagent (Invitrogen) following the manufacturer’s instructions. The extracted RNA concentration and quality were checked using NanoDrop One/OneC (Thermo Scientific). The cDNA was constructed from total RNA using a high-capacity reverse transcriptase kit (Applied Biosystems) following the instructions. Power-Up SYBR green master mix kit (Applied Biosystems) was used to perform RT-PCR analysis. The reaction was performed using a Quant Studio 3 (Applied Biosystems) thermocycler. Specific primers targeting the M-gene of the BCoV (forward primer 5′-CTGGAAGTTGGTGGAGTT-3′ and reverse primer 5′-ATTATCGGCCTAACATACATC-3′) were used to perform RT-PCR (40). Primers for β-actin of bovine (forward primer 5′-CAAGTACCCCATTGAGCACG-3′ and reverse primer 5′-GTCATCTTCTCACGGTTGGC-3′) were used to normalize the viral expression. Tissue samples negative for BCoV were used as a control. The genomic viral expression was normalized following the 2−ΔΔCt method (41).

### 2.8. Confirmation of BCoV by Next Generation Sequencing (NGS)

The samples having high BCoV viral load were subjected to NGS. Total RNA samples were treated with DNase I (New England Biolabs) following the manufacturer’s instructions. The NGS was performed by Azenta Life Sciences (South Plainfield, New Jersey) as described previously (9, 10). Briefly, RNA libraries were constructed using the NEB-Next Ultra II RNA Library Preparation Kit (New England Biolabs), followed by sequencing with Nova-Seq X Plus. The raw sequences were trimmed using Trimmomatic V.0.36 software and mapped with the reference BCoV complete genome sequences. BCoV Mebus isolate (Accession No. BCU00735), and BCoV MARC/2021/01 USA/Nebraska 2021 (Accession No. OP037394) obtained from NCBI were used as reference isolates. The resultant complete genome sequences of BCoV isolates (C1, C2, C5, and C6) were submitted to GenBank and received accession numbers (PX487295, PX486811, and PX485609), respectively.

### 2.9. Multiple Sequence Alignment (MSA) and Phylogenetic Analysis

Multiple sequence alignments and phylogenetic trees were constructed for the complete BCoV genome, S, N, HE, M, and 32 kDa. A Total of 50 BCoV isolates, including four isolates found in this study, were used for MSA analysis. All the reference sequences were obtained from the NCBI database (Supplementary Table 1). The isolates were selected based on BCoV type (enteric/respiratory), geographical location, time of isolation, and genotype. The MSA was performed using Geneious Prime V.11. The phylogenetic trees were constructed using maximum likelihood and the Tamura-Nei model (42), with bootstrap (1,000) replicates. All the phylogenetic trees were constructed using MEGA 12 software (43). Human Coronavirus OC43 (HCoV-O43) isolate LRTI/238/2011 (accession No. KX344031) was used as an out-group in all MSA and phylogenetic analysis.

### 2.10. Structure prediction of key proteins of BCoV isolates

Protein structural prediction for BCoV spike, nucleocapsid, HE, and membrane protein was performed using AlphaFold3 Server (44) using default settings. Amino acid sequences of BCoV key proteins were obtained from NCBI and inserted into the AlphaFold3 Server, and the amino acids showing substitutions compared to SA2 and USDA reference strains were identified. The structure with the highest confidence was visualized using the built-in PyMOL/ChimeraX and used for figure preparation.

### 2.11. BCoV histopathology and immunohistochemistry examination

Bovine tissues submitted to the ISU VDL for diagnosis of respiratory disease were selected for histopathological and IHC analysis. Lung tissue collected at the time of postmortem examination and placed in 10% formalin was processed for routine hematoxylin and eosin (H&E) staining per the ISU VDL standard operating procedures. Lung tissue BCoV RT-PCR positive and included in a diagnosis of BCoV respiratory disease were selected as representative examples of histopathology and IHC detection of BCoV antigen. The IHC performed on the same H&E sections of lungs representing BCoV lesions was conducted on fixed tissue as previously described at the ISU VDL, and per standard operating procedures (5).

### 2.13. Statistical Analysis

The BCoV genomic viral load confirmation and comparison were analyzed using one-way analysis of variance (ANOVA). The statistical analysis was performed using GraphPad Prism software V.9. The P-value less than 0.05 was considered significant, indicated by (* p<0.05, **P < 0.01, and ****P < 0.0001). Data management, descriptive statistics, and regression analysis were conducted using R software (Version 4.1.2; R Foundation for Statistical Computing, Vienna, Austria) within RStudio. Statistical significance was indicated by p-value ≤0.05.

## 3. Results

### 3.1. Descriptive Statistics

The study analyzed a total of 4,505 respiratory samples, and 693 (15.38%) were BCoV RT-PCR positive. The proportion of positive test results varied across different age groups. In the 0-40-day age group, 20.0% were positive (215/1,088). Among the 41–140-day age group, 17.0% were positive (188/1,129). In the 141–240-day age group, 15.0% were positive (195/1,315). In the 241+ day age group, 10.0% were positive (95/973) (Figure 1).

**Figure 1.**
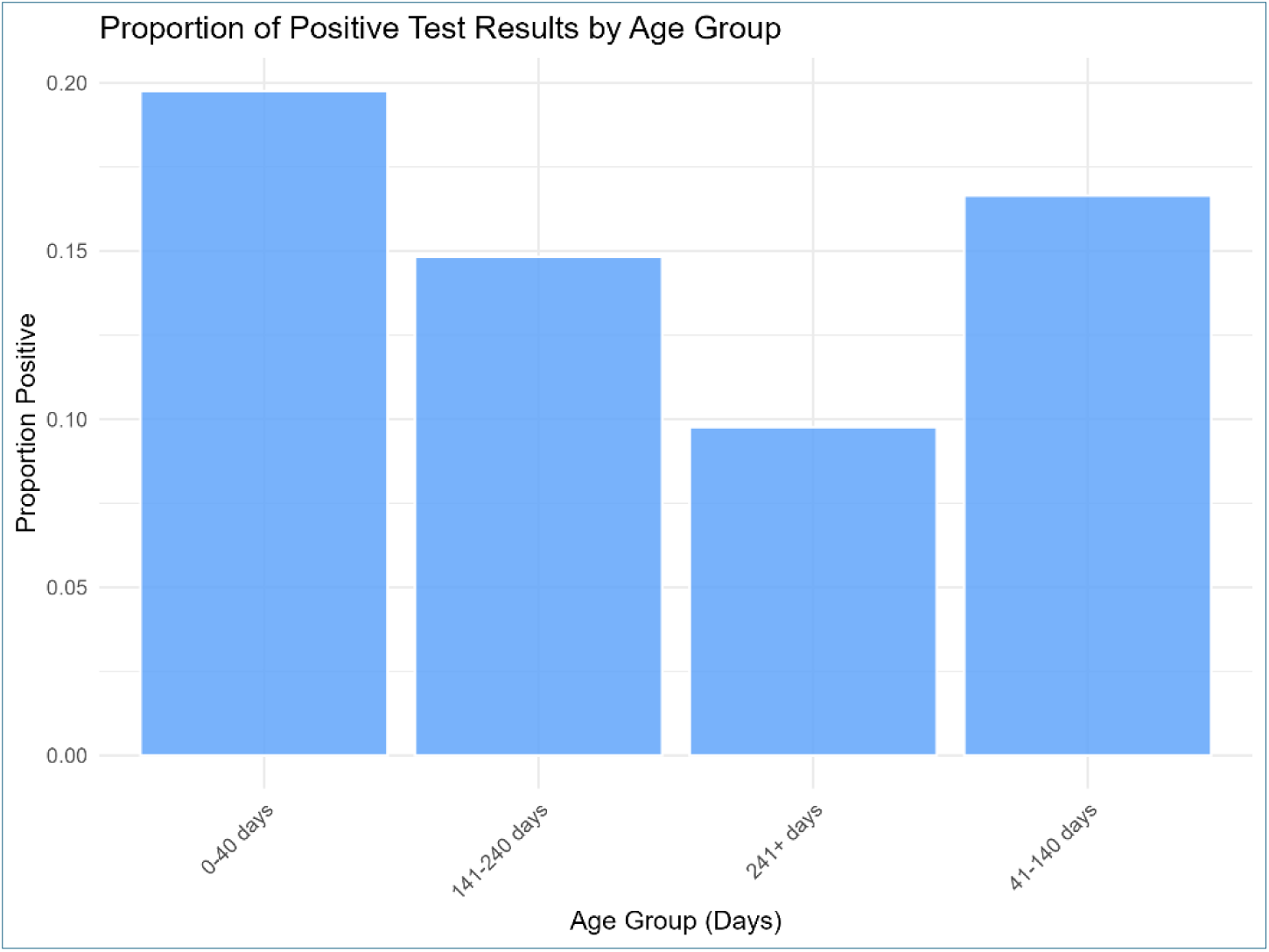
Bovine coronavirus positivity by age group in respiratory tract samples from cattle tested by RT-PCR at the ISU Veterinary Diagnostic Laboratory. This bar chart depicts the proportion of positive test results for bovine coronavirus across four age groups: 0–40 days, 41–140 days, 141–240 days, and 241+ days. The proportion of positive results was highest in the 0–40-day age group.

The proportion of positive RT-PCR by state with at least 100 submissions varied. Ohio had the highest proportion of BCoV-positive samples, with 19.65% (34/173). Indiana followed with 17.05% positive (22/129). Illinois had a positive rate of 16.00% (52/325). Minnesota had 15.38% positive (42/273). Iowa had 14.83% (381/2,569) samples test positive, while Nebraska had the lowest proportion at 14.97% (25/167) positive.

Dairy farms had the highest proportion of positive BCoV samples at 17.65% (57/323). Cow/calf operations followed with 15.81% positive (142/898). Other farm types tested 1,819 samples, with 284 positives, yielding a proportion of 15.61%. Feedlots had the lowest proportion of RT-PCR-positive animals at 14.33% (210/1,465).

### 3.2. Logistic Regression Analysis

A logistic regression model was fitted to examine the relationship between age and the likelihood of testing positive for the condition. As the age in days increased, the odds of test positivity decreased. For each additional day of age, the odds of testing positive decrease by approximately 0.15% (Odds ratio = 0.9985, 95% CI: 0.99797–0.99904, p<0.001), suggesting a negative association between age and test positivity (Figure 2).

**Figure 2.**
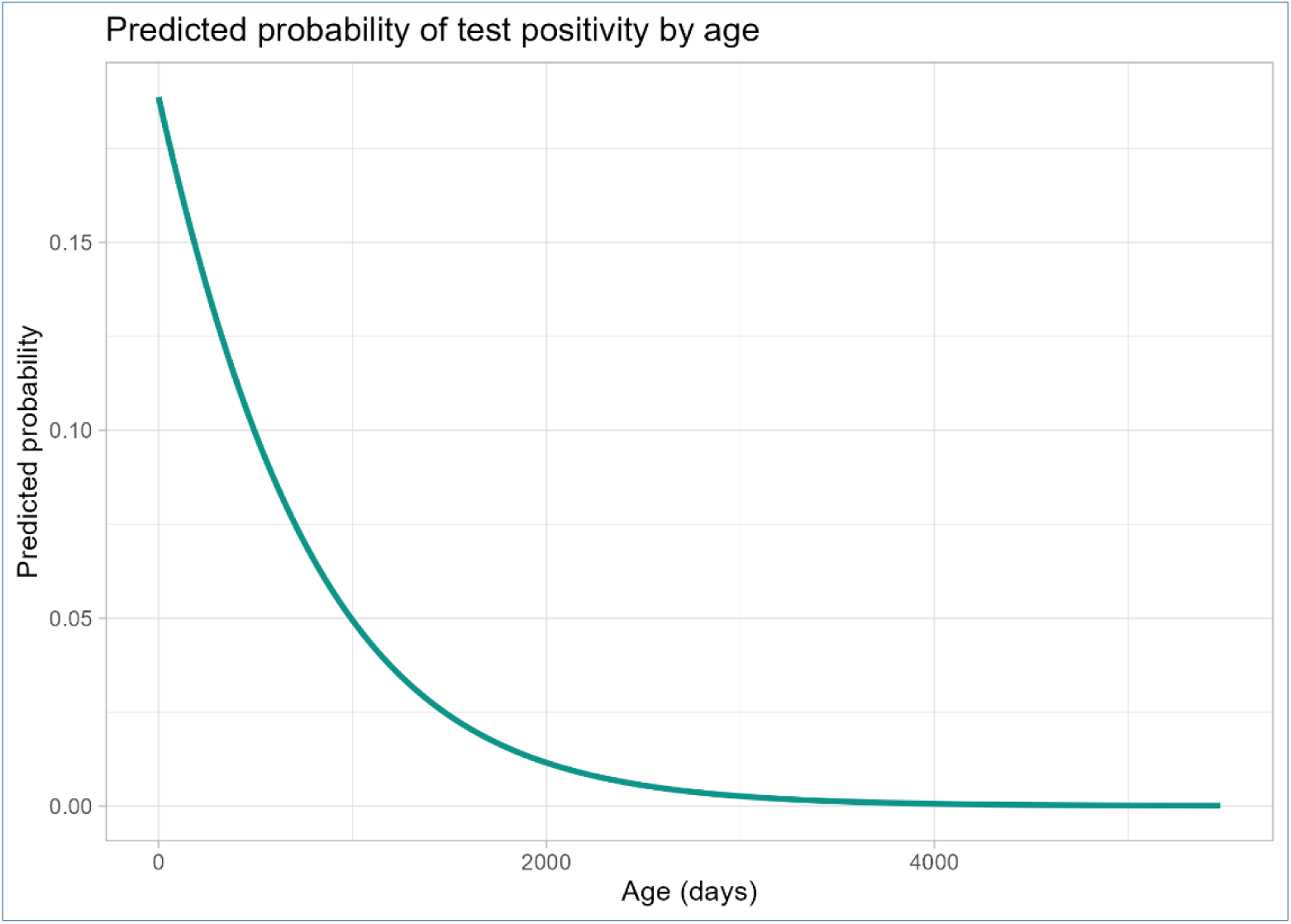
Predicted probability of BCoV test positivity by age from a logistic regression model. The predicted probability of testing positive for BCoV based on age in months was calculated using a logistic regression model. The green line in the figure represents the predicted probability of testing positive for BCoV as a function of age. The x-axis represents age in months, while the y-axis shows the predicted probability of test positivity. As age increases, the predicted probability of testing positive decreases sharply, indicating that older individuals have a lower likelihood of testing positive for BCoV.

The predicted probability of testing positive for BCoV was calculated for each observed age. As age increased, the predicted probability of testing positive decreased, consistent with the regression results.

### 3.3. Trend Analysis

To explore temporal trends, we conducted a trend analysis of the test results over time using LOESS smoothing and the Mann-Kendall (MK) Test (**Figure 3**).

**Figure 3.**
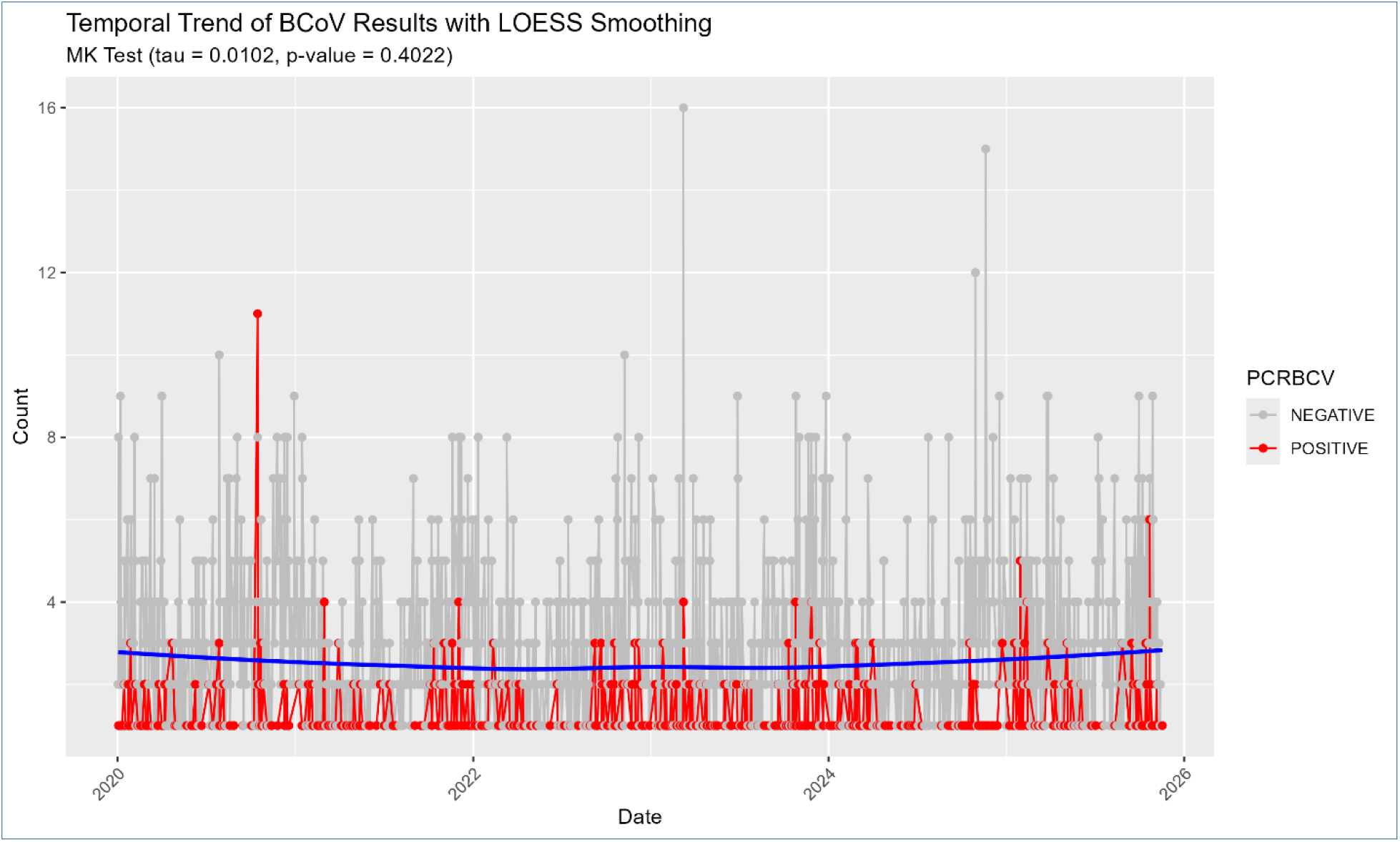
Temporal trends in BCoV test results over the study period, Jan 2020 – Nov 2025. Temporal trends in the count of BCoV positive and negative test results were analyzed using LOESS smoothing and the Mann-Kendall (MK) test. The LOESS smoothing line (blue) reveals fluctuations in the count of positive and negative tests over time, with no significant increases or decreases observed. This pattern is supported by the Mann-Kendall test for monotonic trends (tau = 0.0102, p = 0.4022), indicating no significant monotonic trend in the data over time.

**Figure 4.**
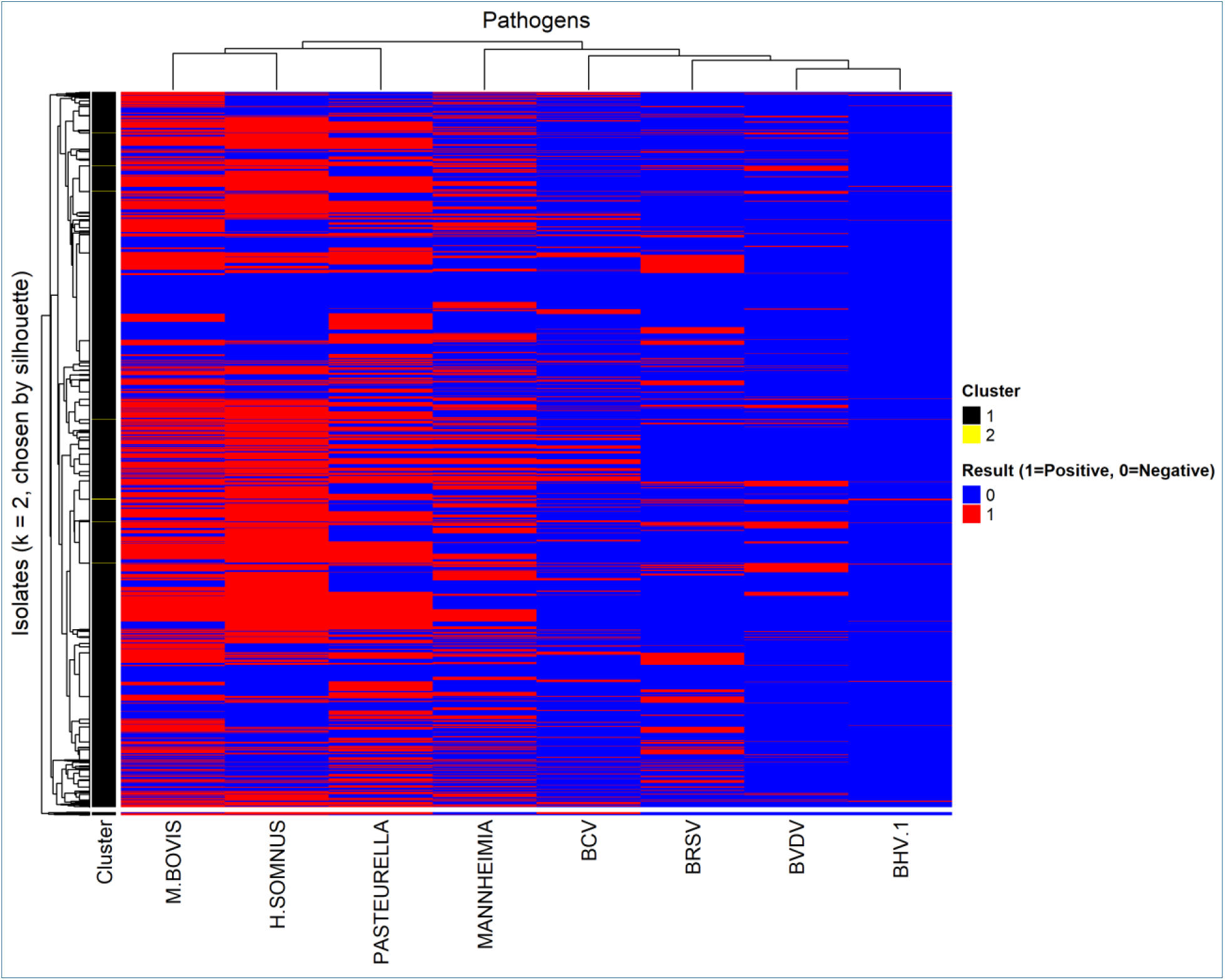
Heatmap of co-infection patterns for bovine bacterial and viral respiratory pathogens. This heatmap illustrates the co-infection patterns of bovine pathogens across two clusters, with the presence (red) or absence (blue) of pathogens indicated for each isolate. The columns represent the pathogens, and the rows represent the isolates, including both bacterial and viral pathogens.

The LOESS smoothing line showed fluctuation in the count of BCoV positive and negative tests over time. It appeared relatively stable with no main variations in increases or decreases over time. This observation was further supported by the results of the MK test for monotonic trends, which yielded a tau value of 0.0102 (p = 0.4022). This result indicates that there was no significant monotonic trend in the data over time. The positive value suggests a very slight upward trend, but the *p*-value indicates that this trend is not statistically significant.

### 3.4. Assessing the co-infections of BCoV and other viral and bacterial pathogens

Our results show the co-infection patterns of bovine pathogens across two clusters, with the presence (red) or absence (blue) of pathogens indicated for each isolate. The columns represent the pathogens, including both bacterial and viral pathogens. The majority of bacterial pathogens, *H. somnus, M. bovis, M. haemolytica*, and *P. multocida*, are more prominently represented in the samples, with higher proportions of RT-PCR positive results. In contrast, viral pathogens (BCV, BHV-1, BRSV, BVDV) are less frequently detected.

Co-occurrence patterns (columns in the heatmap) are evident. Among the viral pathogens, BHV-1 co-occurs most frequently with BVDV, followed by BRSV and BCoV. Among the bacterial pathogens, *H. somnus* co-occurs most often with *M. bovis*, followed by *P. multocida* and *M. haemolytica*. These patterns suggest that, within this dataset, viral pathogens have a higher likelihood of co-occurrence with other viral pathogens, and bacterial pathogens tend to co-occur more frequently with other bacterial pathogens, with little to no co-occurrence between viral and bacterial pathogens. However, these co-occurrence patterns might be influenced by the stage of the disease process at the time of sample collection, since these are field samples and do not represent controlled experimental infections.

When analyzing the co-infection profiles of isolates (rows of the heatmap), two clusters were identified. Cluster 1, which comprised most of the samples (99.55%), was characterized by higher proportions of bacterial pathogens, including *H. somnus* (56.12%), *M. bovis* (57.58%), *P. multocida* (50.89%), and *M. haemolytica* (36.84%). In contrast, viral pathogens were less prevalent in Cluster 1, with BCoV present in only 14.99%, BHV-1 in 1.53%, BRSV in 18.20%, and BVDV in 10.13%.

In contrast, Cluster 2, containing just 0.45% of the samples, exhibited high proportions of both viral and bacterial pathogens. BCoV was detected in 94.44% of the samples, BHV-1 in 66.67%, and BRSV in 72.22%. This cluster also showed a high prevalence of bacterial pathogens, with *M. haemolytica* detected in 77.78% and *P. multocida* in 72.22% of samples, *M. bovis* in 57.58%, and *H. somnus* in 56.12% of samples.

### 3.5. Detection of Respiratory BCoV infection in bovine lung tissues

Bovine lung tissue samples were collected from 13 different farms. The samples named B1 – B7 were collected in 2023, and C1 – C6 were collected in 2024. All the samples were tested for BCoV infection by RT-PCR. Results showed the presence of BCoV viral genome with lower viral load in samples B1, B2, B5, C3, and C4 (Figure 5). While high BCoV viral genome copy number was observed in lung samples collected from C1, C2, C5, and C6 (Figure 5). Therefore, these four samples were further processed for the BCoV genome sequencing.

**Figure 5:**
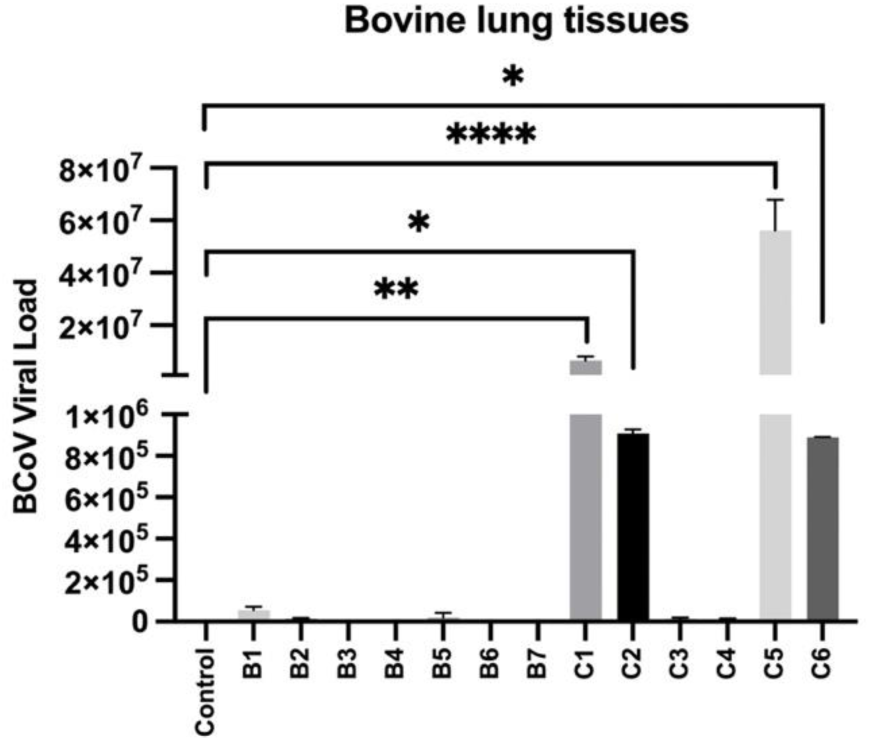
BCoV genomic viral load confirmation by RT-PCR. Bovine lung samples from thirteen infected animals were tested for BCoV viral load. A tissue sample negative for BCoV was used as a control.

### 3.6. Complete genome sequencing of BCoV-C1, C2, C5, and C6 on clinical samples

The complete genome sequences of BCoV were obtained from lung tissue C1, C2, C5, and C6 by NGS analysis. The sequences were named after their respective groups, assembled, and deposited in the NCBI GenBank database under accession numbers BCoV-C1 (PX485608), BCoV-C2 (PX487295), BCoV-C5 (PX486811), and BCoV-C6 (PX485609). The complete genomic size of BCoV-C1 and BCoV-C5 were 31,006 nucleotides, whereas those of BCoV-C2 and BCoV-C6 were 31,005 nucleotides. Since these sequences were retrieved from bovine lungs and were considered BCoV respiratory samples, we aligned these sequences with the reference BCoV respiratory isolate MARC/2021/01 from Nebraska, USA, and the BCoV reference Mebus isolate. BCoV-C1 and BCoV-C2 showed 99.7% similarity, BCoV-C5 showed 99.5% similarity, and BCoV-C6 showed 99.9% similarity with the reference MARC/2021/01 isolate. In contrast, the BCoV reference showed 98.2% similarity between the Mebus isolates BCoV-C1 and BCoV-C6, and 98.1% similarity between BCoV-C2 and BCoV-C5. The viral genome of all four BCoV obtained in this study was organized with the 5’UTR region followed by ORF1ab, which occupied two-thirds of the genome (Table 1, 2). The remaining one-third of the genome contains five main structural proteins (HE, S, E, M, and N) with internal proteins in between nucleocapsid (Table 1, 2). This region also contains four non-structural proteins (NSP) (32 kDa, 4.9 kDa, 4.8 kDa, and 12.7 kDa) (Table 1, 2). The BCoV-C2 sequences have a small 4.8 kDa NSP and an extra protein of 87 nucleotide size at the same region, which was designated protein X (Table 1).

**Table 1:** Genome organization of BCoV-C1 and BCoV-C2 whole genome sequences.

| Gene | BCoV-C1 |  |  |  | BCoV-C2 |  |  |  |
| --- | --- | --- | --- | --- | --- | --- | --- | --- |
|  | Start | Stop | Size (nt) | Size (aa) | Start | Stop | Size (nt) | Size (aa) |
| 5'UTR | 1 | 200 | 200 | - | 1 | 200 | 200 | - |
| ORF1a | 201 | 13352 | 13152 | 4384 | 201 | 13352 | 13152 | 4384 |
| ORF1ab | 201 | 21484 | 21284 | 7094 | 201 | 21484 | 21284 | 7094 |
| 32 kDa | 21494 | 22330 | 837 | 279 | 21494 | 22330 | 837 | 279 |
| Hemagglutinin-Esterase | 22342 | 23616 | 1275 | 425 | 22342 | 23616 | 1275 | 425 |
| Spike | 23631 | 27722 | 4092 | 1364 | 23631 | 27722 | 4092 | 1364 |
| 4.9 kDa | 27712 | 27801 | 90 | 30 | 27712 | 27801 | 90 | 30 |
| 4.8 kDa | 27879 | 28004 | 126 | 42 | 27879 | 27944 | 66 | 22 |
| Protein X | - | - | - | - | 27994 | 28080 | 87 | 29 |
| 12.7 kDa | 28084 | 28413 | 330 | 110 | 28083 | 28412 | 330 | 110 |
| Envelope | 28400 | 28654 | 255 | 85 | 28399 | 28653 | 255 | 85 |
| Membrane | 28669 | 29361 | 693 | 231 | 28668 | 29360 | 693 | 231 |
| Nucleocapsid | 29371 | 30717 | 1347 | 449 | 29370 | 30716 | 1347 | 449 |
| Internal | 29432 | 30055 | 624 | 208 | 29431 | 30054 | 624 | 208 |
| 3'UTR | 30718 | 31006 | 289 | - | 30717 | 31005 | 289 | - |

**Table 2:** Genome organization of BCoV-C5 and BCoV-C6 whole genome sequences.

| Gene | BCoV-C5 |  |  |  | BCoV-C6 |  |  |  |
| --- | --- | --- | --- | --- | --- | --- | --- | --- |
|  | Start | Stop | Size (nt) | Size (aa) | Start | Stop | Size (nt) | Size (aa) |
| 5'UTR | 1 | 200 | 200 | - | 1 | 200 | 200 | - |
| ORF1a | 201 | 13352 | 13152 | 4384 | 201 | 13352 | 13152 | 4384 |
| ORF1ab | 201 | 21484 | 21284 | 7094 | 201 | 21484 | 21284 | 7094 |
| 32 kDa | 21494 | 22330 | 837 | 279 | 21494 | 22330 | 837 | 279 |
| Hemagglutinin-Esterase | 22342 | 23616 | 1275 | 425 | 22342 | 23616 | 1275 | 425 |
| Spike | 23631 | 27722 | 4092 | 1364 | 23631 | 27722 | 4092 | 1364 |
| 4.9 kDa | 27712 | 27801 | 90 | 30 | 27712 | 27801 | 90 | 30 |
| 4.8 kDa | 27879 | 28004 | 126 | 42 | 27879 | 27956 | 78 | 26 |
| 12.7 kDa | 28084 | 28413 | 330 | 110 | 28083 | 28412 | 330 | 110 |
| Envelope | 28400 | 28654 | 255 | 85 | 28399 | 28653 | 255 | 85 |
| Membrane | 28669 | 29361 | 693 | 231 | 28668 | 29360 | 693 | 231 |
| Nucleocapsid | 29371 | 30717 | 1347 | 449 | 29370 | 30716 | 1347 | 449 |
| Internal | 29432 | 30055 | 624 | 208 | 29431 | 30054 | 624 | 208 |
| 3'UTR | 30718 | 31006 | 289 | - | 30717 | 31005 | 289 | - |

### 3.7. Phylogenetic analysis of BCoV-C1, C2, C5, and C6 whole genomes in comparison with BCoV reference sequences

A phylogenetic tree was constructed using 48 complete genome sequences, including BCoV-C1, C2, C5, and C6, to determine the genotype group reported in this study. The phylogenetic tree was divided into four groups (GIa, GIb, GIIa, and GIIb) based on established BCoV genotypes. The BCoV-C1, C2, C5, and C6 isolates clustered with BCoV respiratory isolate MARC/2021/01 and enteric isolate VDC/2022/06, both from Nebraska, within group GIIb (Figure 6). Notably, most American BCoV isolates from Nebraska, Iowa, Oregon, Kansas, Ohio, and Pennsylvania collected between 2014 and 2024 also belong to group GIIb. Some BCoV sequences from Ohio in 1998 and 2001, clustered with BCoV isolates from Japan and China, were assigned to group GIIa. BCoV isolates from Ireland, Turkey, and France grouped within genotype GIb. Whereas, the classical BCoV isolates like Mebus, Quebec, Kakegawa, and the BCoV-13 isolate recently characterized by our lab clustered within genotype group GIa (Figure 6).

**Figure 6:**
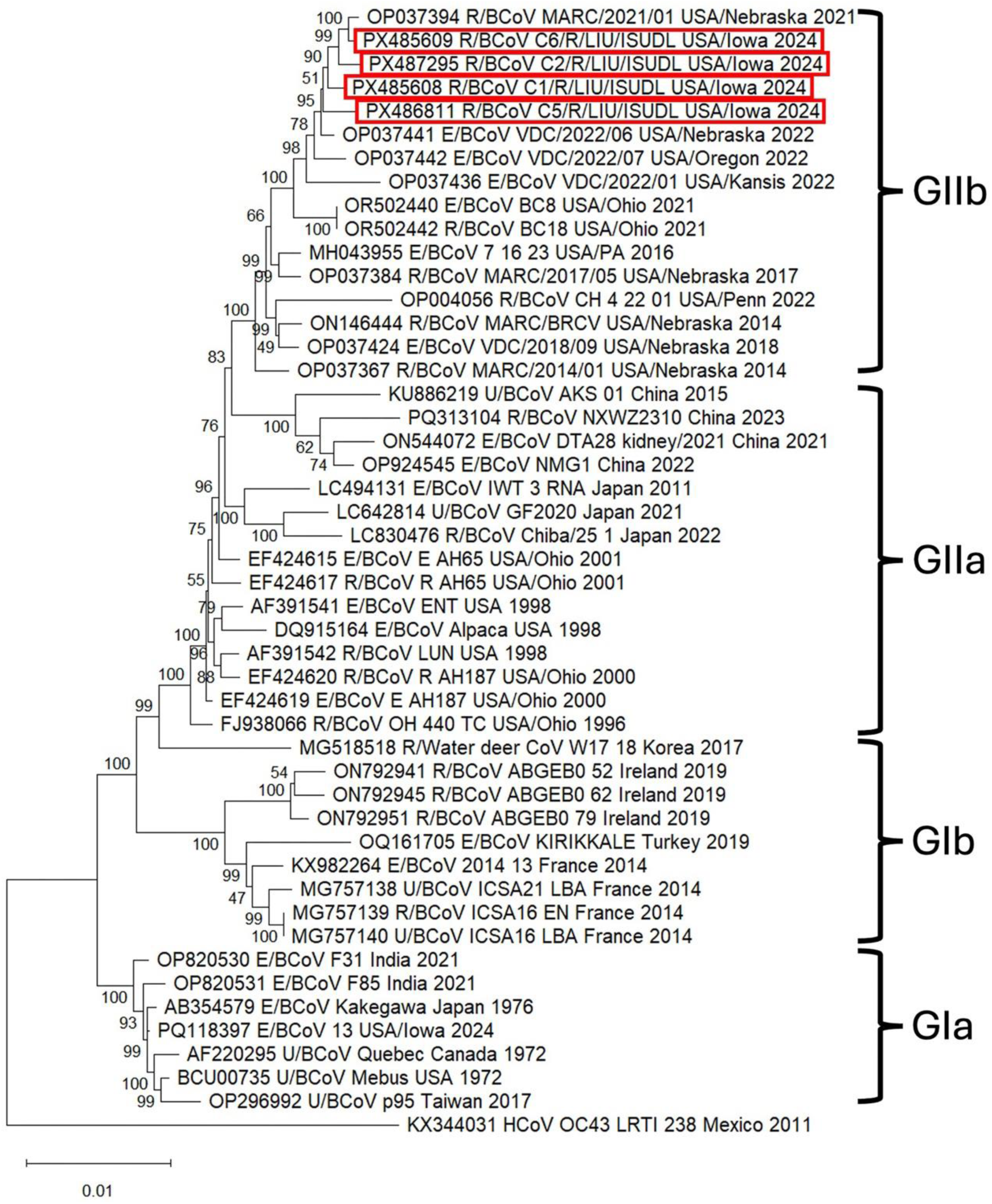
Phylogenetic tree of 48 BCoV complete genome sequences. The phylogenetic tree was constructed of 44 BCoV complete genomes selected from GenBank and included BCoV-C1, C2, C5, and C6 whole genomes from this study. The maximum log likelihood of the tree was -84,232.27, associated with the taxa clustered together (1,000 replicates), shown at each branch. The sequences generated in this study are shown in red boxes. BCoV groups are indicated at the right side of the tree. E represents enteric, R represents respiratory, and U represents an unknown type of BCoV. Human CoV OC43 was used as an outgroup. The tree was generated using MEGA12 software.

### 3.8. Phylogenetic analysis of Hemagglutinin-Esterase (HE) protein of BCoV-C1, C2, C5, and C6

The phylogenetic analysis of the HE protein was conducted using 50 BCoV sequences, including the BCoV-C1, C2, C5, and C6 reported in this study. The reference BCoV enteric and respiratory strains were selected based on geographical location, isolation time, and genotypes (Supplementary Table 1). The phylogenetic analysis of the HE protein showed that BCoV-C1, C2, C5, and C6 clustered together with other American BCoV enteric and respiratory strains from Ohio and Nebraska (Figure 7A). To further elaborate on the variation in HE protein of BCoV-C1, C2, C5, and C6, we performed multiple sequence alignment among these 50 BCoV sequences (Figure 7B). There were five novel amino acid substitutions discovered in BCoV-C2, C5, and C6 that were completely different from those observed in all reference sequences (Figure 7B – 7G). Among these substitutions, three were reported in BCoV-C2, while one each was found in BCoV-C5 and BCoV-C6. The first substitution occurred in the HE protein of BCoV-C2, where leucine was replaced by arginine at position 235 (L235R) (Figure 7B, 7D). The second substitution was observed at position 337 of the HE protein in BCoV-C2, where threonine was encoded instead of isoleucine (I337T) present in the other 49 reference sequences (Figure 7B, 7E). The third substitution is at position 375 of the HE protein of BCoV-C2, where valine replaces alanine (A375V) (Figure 7B, 7G). The fourth substitution was observed at position 349 of the HE protein, where BCoV-C5, BCoV-C6, and VDC/06 isolates from Nebraska replace serine with alanine (S349A) (Figure 7B, 7F).

**Figure 7:**
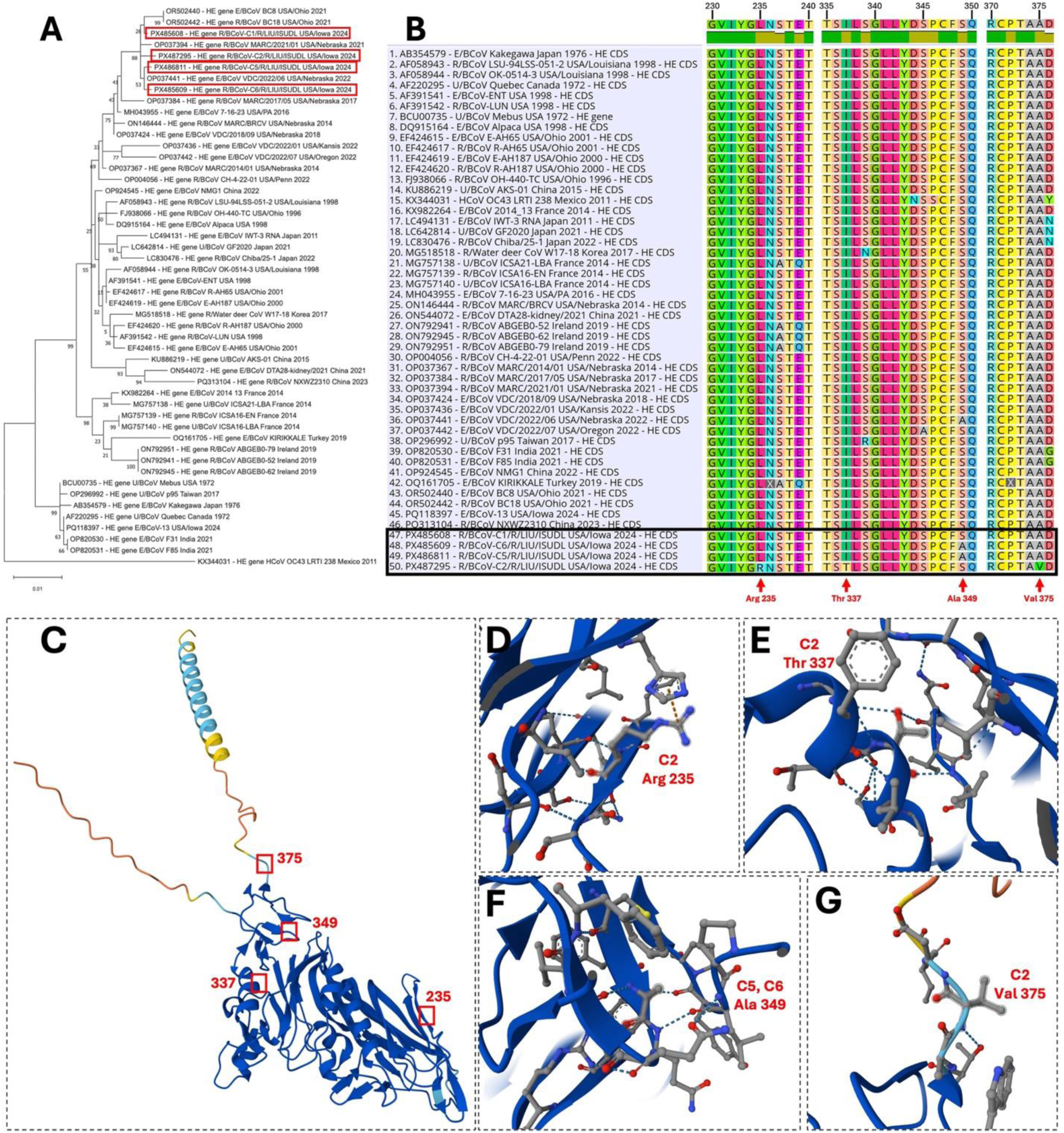
Phylogenetic tree construction and divergence analysis of HE protein. (A) The phylogenetic tree was constructed of the HE gene of 50 BCoV, including BCoV-C1, C2, C5, and C6. The maximum log likelihood of the tree was -3,738.11, associated with the taxa clustered together (1,000 replicates), shown at each branch. The gene sequences generated in this study are shown in red boxes. The tree was generated using MEGA12 software. (B) Multiple sequence alignment (MSA) of the HE protein of 50 BCoV was performed using Geneious Prime V.11. The amino acid sequences generated in this study are shown in a black box. Mutations are indicated with a red arrowhead at the bottom. (C) Protein structure of HE BCoV-C2. The mutations are indicated with a red box and numbering. (D) Protein structure of HE of BCoV-C2 showing arginine at position 235; (E) threonine at position 337. (F) Protein structure of HE of BCoV-C5 and BCoV-C6 showing alanine at position 349. (G) The HE protein of BCoV-C2 shows valine at position 375. The protein structure was designed using AlphaFold Server 3.

### 3.9. Notable substitutions in the spike glycoprotein of BCoV-C1, C2, C5, and C6

A phylogenetic tree was constructed based on the spike glycoprotein sequences of 50 BCoV, including BCoV-C1, C2, C5, and C6 whole genomes generated in this study. Results showed that the spike glycoprotein of BCoV-C1, C2, C5, and C6 clustered with the enteric American BCoV strain from Oregon (VDC/2022/07) and both enteric and respiratory strains from Nebraska (VDC/2022/06 and MARC/2021/01) (Figure 8A). To identify notable substitutions in the spike glycoprotein, multiple sequence alignment was performed among the 50 BCoV sequences (Figure 9A). A total of sixteen novel amino acid substitutions were identified in BCoV-C1, C2, C5, and C6 that were completely different from those observed in all other 46 reference strains (Figure 9A). Among these substitutions, three were reported in BCoV-C1, five in BCoV-C2, six in BCoV-C5, and two in BCoV-C6 (Figure 9B – 9D). The substitution in the spike glycoprotein of BCoV-C1 was alanine replaced with serine at position 273 (A273S), and threonine was replaced with serine at position 772 (T772S) (Figure 9B). In the spike glycoprotein of BCoV-C2, methionine was replaced by threonine at position 240 (T240M), threonine by isoleucine at position 617 (T617I), lysine by glutamic acid at position 621 (K621E), lysine by arginine at position 1170 (K1170R), and aspartic acid by alanine at position 1296 (D1296A) (Figure 9C). In the spike glycoprotein of BCoV-C5; leucine was replaced by phenylalanine at position 232 (F232L), histidine by tyrosine at position 244 (H244Y), alanine by serine at position 435 (A435S), threonine by isoleucine at position 447 (T447I), isoleucine by valine at position 644 (I644V), and valine by phenylalanine at position 744 (V744F) (Figure 9D). Another notable substitution was observed at position 154 of the spike glycoprotein: some strains encoded leucine, some encoded phenylalanine, whereas BCoV-C6 encoded serine (L/F154S), which was distinct from all other strains included in the study. Another substitution reported in the spike glycoprotein of BCoV-C6 was isoleucine replacing methionine at position 680 (M680I) (Figure 9E).

**Figure 8:**
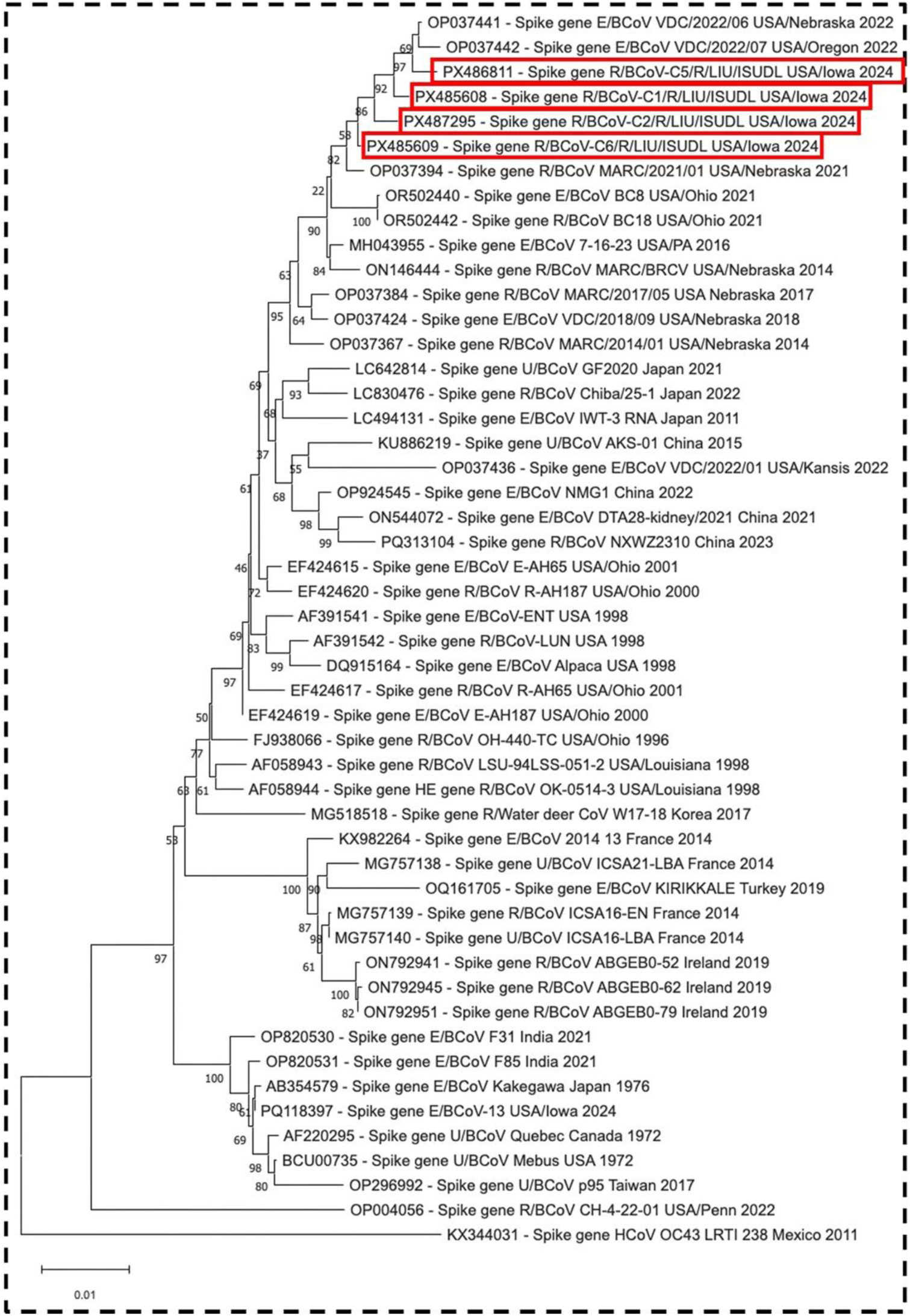
Phylogenetic tree construction of spike glycoprotein. The phylogenetic tree was constructed based on the spike glycoprotein of 50 BCoV sequences, including BCoV-C1, C2, C5, and C6. The maximum log likelihood of the tree was -14,099.70, associated with the taxa clustered together (1,000 replicates), shown at each branch. The spike glycoprotein sequences generated in this study are shown in red boxes. The tree was generated using MEGA12 software.

**Figure 9:**
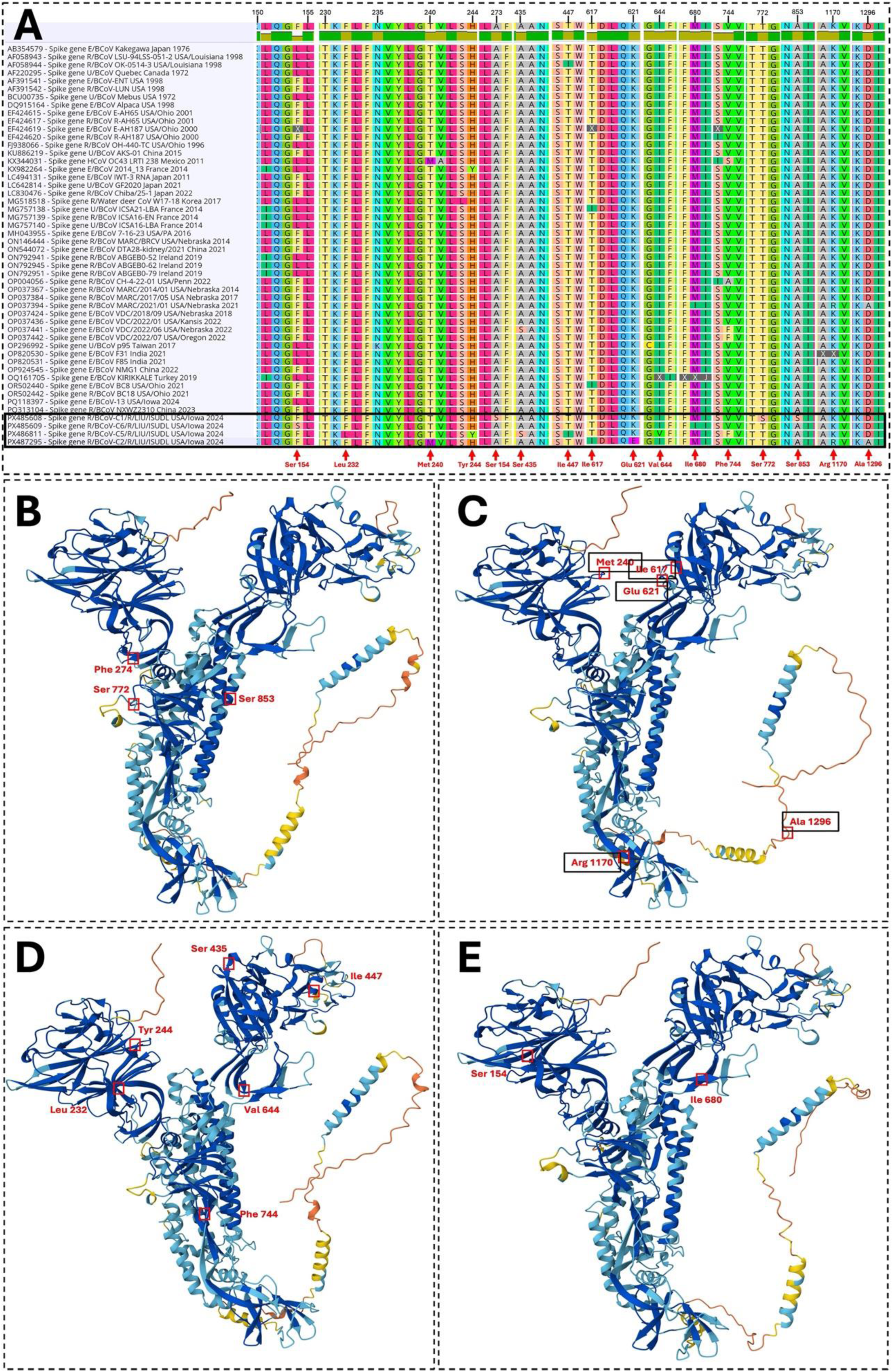
Divergence analysis in spike glycoprotein. (A) MSA of the spike glycoprotein of 50 BCoV was performed using Geneious Prime V.11. The spike gene amino acid sequences generated in this study are shown in a black box. Mutations are indicated with a red arrowhead at the bottom. (B) Protein structure of spike glycoprotein of BCoV-C1; (C) BCoV-C2; (D) BCoV-C5; and (E) BCoV-C6. The mutations are indicated with a red box and numbering. All protein structures were designed using AlphaFold Server 3.

### 3.10. Variations in the Nucleocapsid protein of BCoV-C2, C5, and C6 isolates

A phylogenetic tree was constructed based on the nucleocapsid protein sequences of 50 BCoV, including BCoV-C1, C2, C5, and C6, generated in this study. Results showed that the nucleocapsid protein of BCoV-C1 clustered with the BCoV strains from Ohio (E-AH65/2001) and R-BC18/2021 (Figure 10A). The nucleocapsid protein of BCoV-C2 clustered with respiratory BCoV strains from Nebraska (MARC/2021/01) and enteric strains from China (DTA28/2021) (Figure 10A). The nucleocapsid protein of BCoV-C5 clustered with respiratory BCoV strains from Nebraska (MARC/BRCV/2014) and enteric strains from Oregon (VDC/2022/07) (Figure 10A). The nucleocapsid protein of BCoV-C6 clustered with the enteric BCoV strains from Pennsylvania (7-16-23/2016) (Figure 10A). Multiple sequence alignment was performed to identify notable variations in the amino acid sequences of BCoV-C1, C2, C5, and C6 (Figure 10B). Four novel mutations were observed in the nucleocapsid protein of BCoV-C5, two in BCoV-C2, and one in BCoV-C6 (Figure 10B, 10C, 10D). The substitution in the nucleocapsid protein of BCoV-C2 was arginine replaced with cysteine at position 203 (R203C), and serine replaced with phenylalanine at position 209 (S209F) (Figure 10B, 10C). In the nucleocapsid protein of BCoV-C5, glutamine was replaced by leucine at position 52 (Q52L), glutamine by histidine at position 184 (Q184H), lysine by arginine at position 392 (K392R), and threonine by isoleucine at position 442 (T442I) (Figure 10B, 10D). The only mutation identified in the nucleocapsid protein of BCoV-C6 was the substitution of serine with lysine at position 364 (S364L) (Figure 10B).

**Figure 10:**
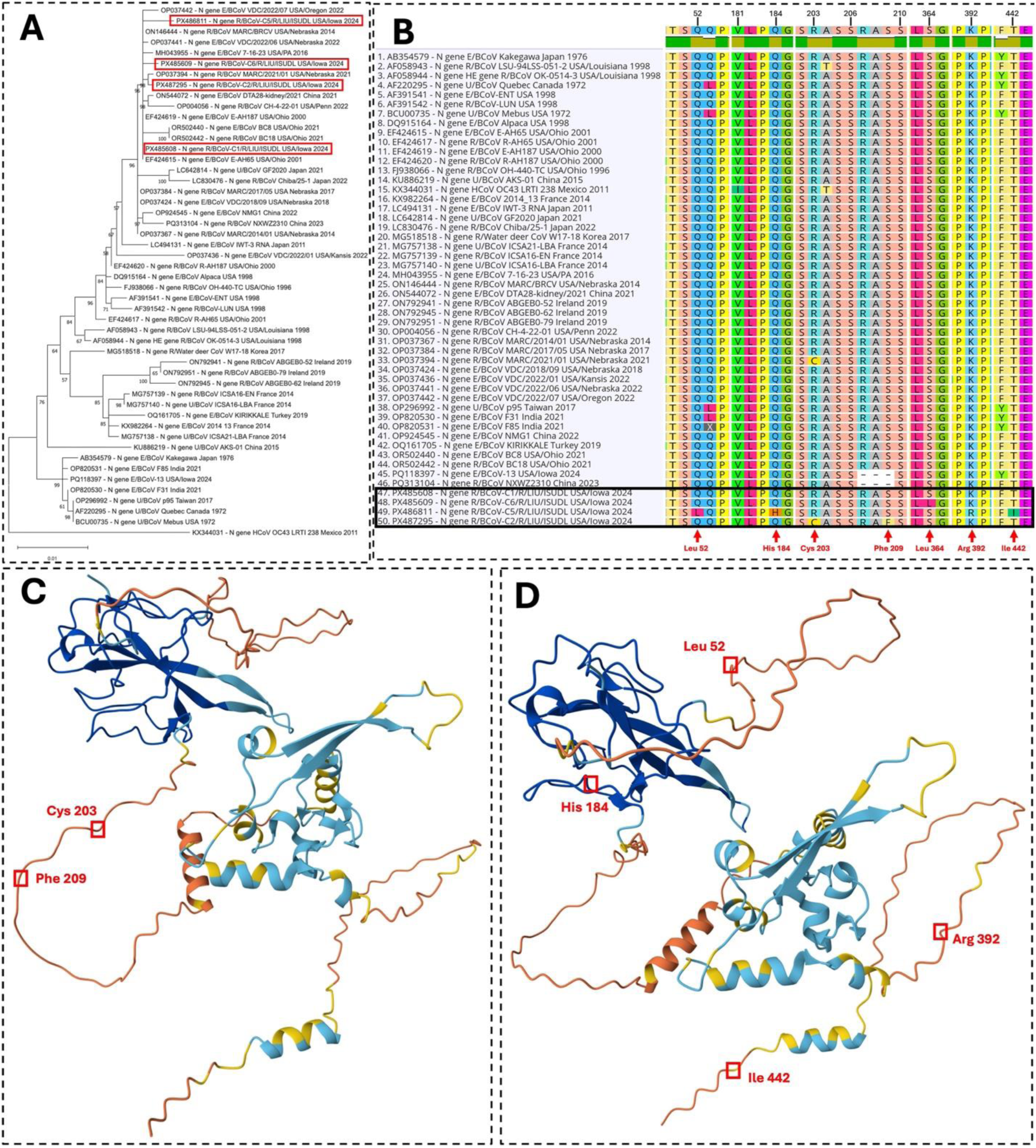
Phylogenetic tree and divergence analysis in the nucleocapsid of BCoV-C1, C2, C5, and C6. (A) The phylogenetic tree was constructed based on the nucleocapsid nucleotide sequences of 50 BCoV, including BCoV-C1, C2, C5, and C6, included in this study. The maximum log likelihood of the tree was -3,456.66, associated with the taxa clustered together (1,000 replicates), shown at each branch. The nucleoprotein gene sequences generated in this study are shown in red boxes. The tree was generated using MEGA12 software. (B) MSA of the nucleocapsid protein sequences of 50 BCoV was performed using Geneious Prime V.11. The spike gene amino acid sequences generated in this study are shown in a black box. Mutations are indicated with a red arrowhead at the bottom. (C) Protein structure of nucleocapsid protein of BCoV-C2, and (D) BCoV-C5. The mutations are indicated with a red box and numbering. All protein structures were designed using AlphaFold Server 3.

### 3.11. Mutations in the membrane protein of the BCoV-C5 isolate

Phylogenetic analysis of the membrane protein showed that BCoV-C1, C5, and C6 clustered with the American sequences VDC/2022/01 from Kansas and VDC/2022/06 from Nebraska (Figure 11A). The BCoV-C2 also clustered with American sequences MARC/2021/01 from Nebraska and VDC/2022/07 from Oregon. Multiple sequence alignment of the membrane protein revealed two novel amino acid substitutions in BCoV-C5 (Figure 11B). The first mutation occurred at position 197 of the membrane protein of BCoV-C5, where serine was replaced by threonine (S197T). The second substitution was observed at position 221, where some isolates encoded methionine or leucine, whereas BCoV-C5 encoded valine (M/L221V) (Figure 11B).

**Figure 11:**
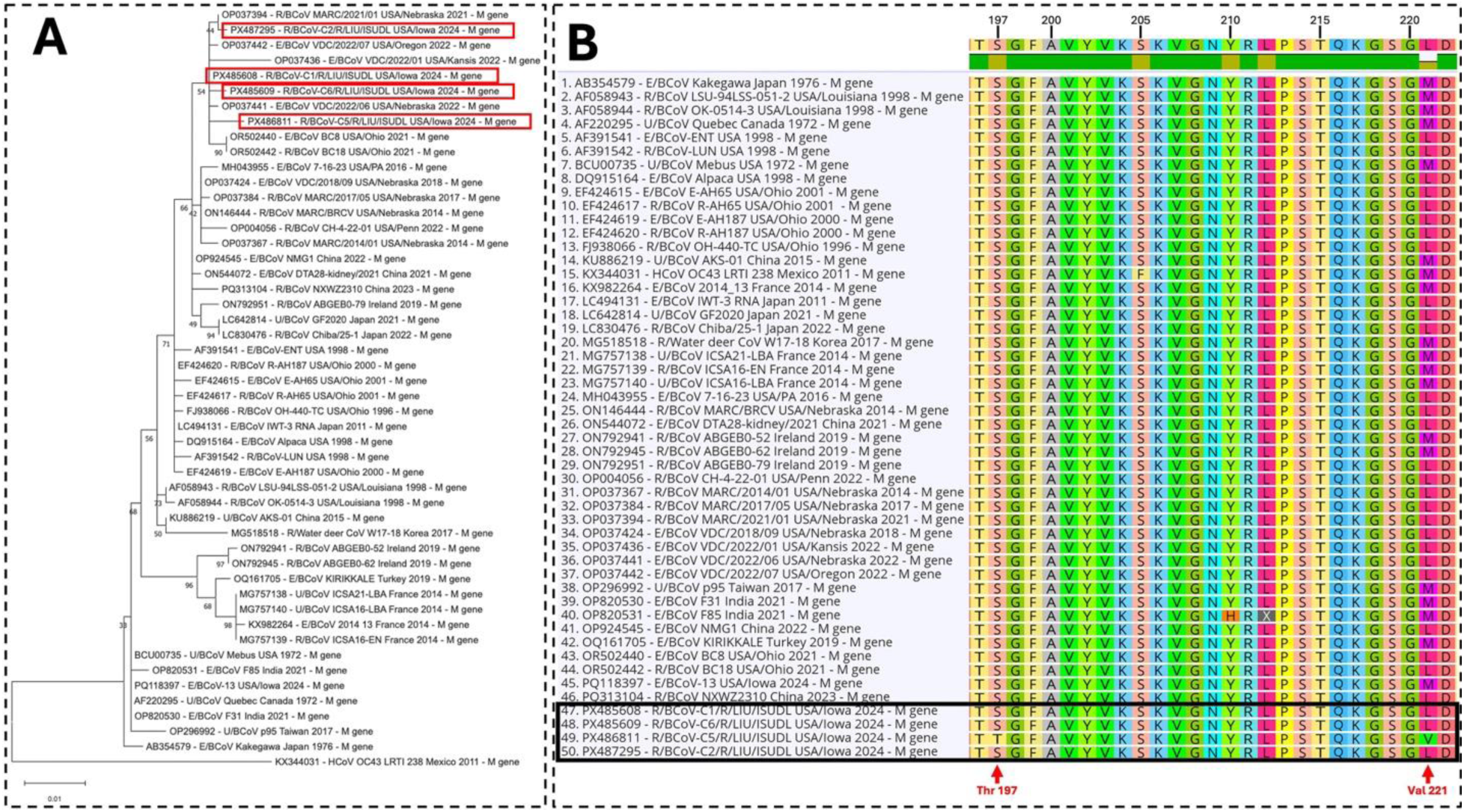
Phylogenetic tree and divergence analysis in the membrane protein of BCoV-C1, C2, C5, and C6. (A) The phylogenetic tree was constructed based on the membrane gene nucleotide sequences of 50 BCoV, including BCoV-C1, C2, C5, and C6, generated in this study. The maximum log likelihood of the tree was -1,838.07, associated with the taxa clustered together (1,000 replicates), shown at each branch. The membrane gene sequences generated in this study are shown in red boxes. The tree was generated using MEGA12 software. (B) MSA of the membrane protein sequences of 50 BCoV was performed using Geneious Prime V.11. The membrane amino acid sequences generated in this study are shown in a black box. Mutations are indicated with a red arrowhead at the bottom.

### 3.12. Representative histopathological lesions induced by the currently circulating BCoV respiratory strains in cattle

BCoV replication occurs in the respiratory tract of bovine species, which may cause histopathological lesions associated with epithelial insult in the trachea and conducting airways of the lung (5). Depending on the stage of the disease process when lung samples were collected, bronchi and bronchiolar epithelium may demonstrate proliferation (subacute; Figure 12A) or attenuation and necrosis (acute; Figure 12C) with inflammation in the airway lumen and surrounding affected lung parenchyma. Immunohistochemistry targeting BCoV antigen, indicating the presence of virus, is consistent with the location of airway lesions in subacute (Figure 12B) and acute (Figure 12D) infections as evidenced by BCoV signals in the cytoplasm of the conducting airway epithelium. The lung lesions in Figures 12A and B were associated with a *M. haemolytica* co-infection, and Figures 12C, and D associated with *Bibersteinia trehalosi* and *P. multocida* co-infections.

**Figure 12.**
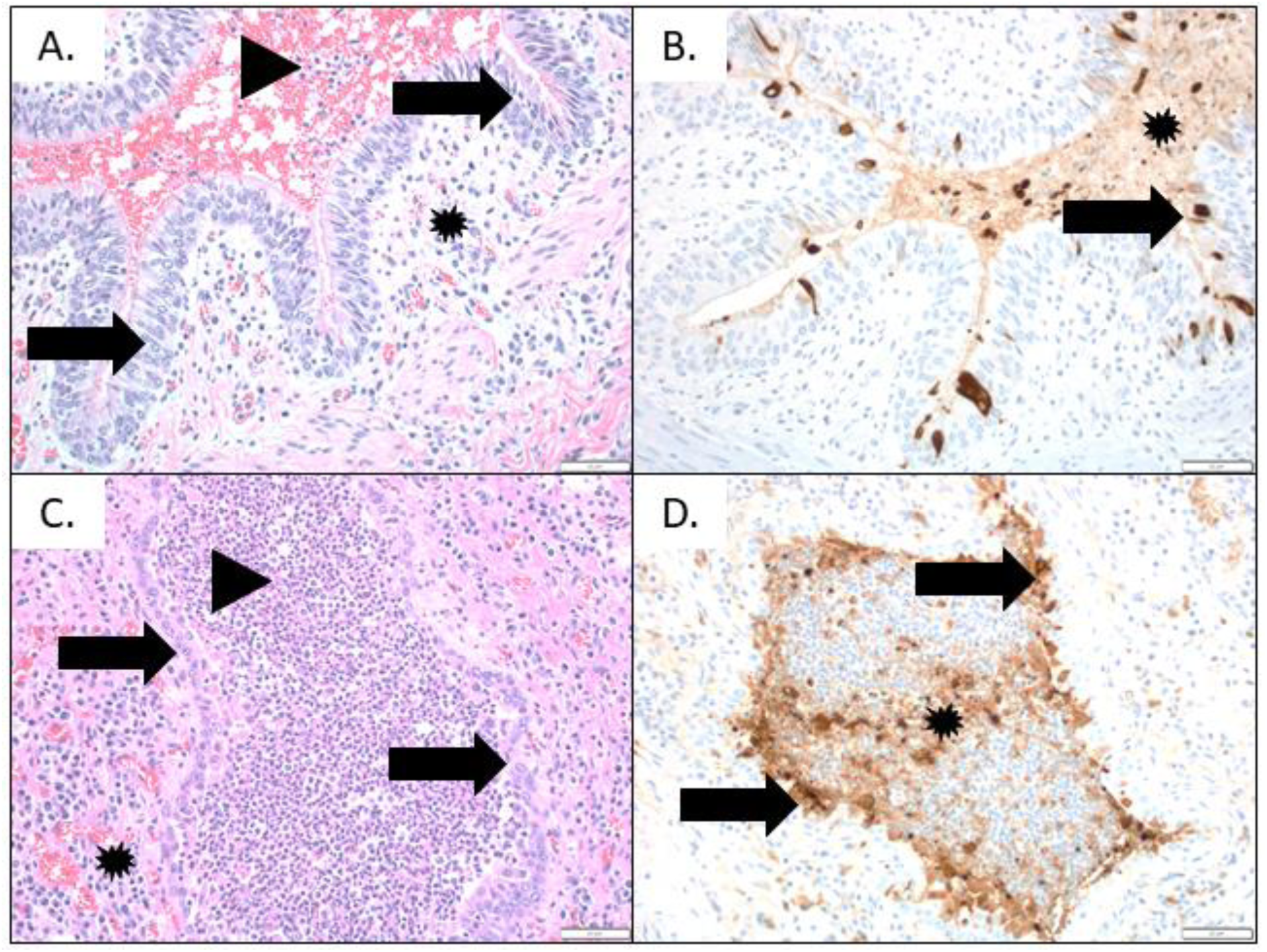
Bovine coronavirus (BCoV) microscopic lung lesions with immunohistochemistry direct detection from two different submissions to the Iowa State University Veterinary Diagnostic Laboratory. (A) Bronchus from a three-week-old calf with respiratory disease. Epithelium is proliferative, stacked or stratified (arrow), and the lumen contains hemorrhage and low numbers of neutrophils (arrowhead). The propria submucosa is infiltrated with low numbers of neutrophils, lymphocytes, and macrophages (asterisk). Adjacent pulmonary parenchyma contains multifocal regions of necrosis and fibrin surrounded by degenerate neutrophils and pleuritis. BCoV reverse transcriptase real-time PCR cycle threshold value 29.1. (200X, H&E) (B) BCoV immunohistochemistry demonstrates cell-associated, immunoreactive brown signals in affected airway epithelium (arrow) and sloughed immunoreactive debris in the lumen (asterisk). (200X, IHC) (C) Bronchiole from a five-month-old calf with respiratory disease consisting of coughing and nasal exudate. Bronchiolar epithelium is attenuated and contains occasional neutrophils (arrow). The airway lumen contains large numbers of neutrophils and macrophages (arrowhead) and mild airway cuffing with lymphocytes and plasma cells (asterisk). Adjacent pulmonary parenchyma is infiltrated with large numbers of degenerate neutrophils and macrophages and type II pneumocyte hyperplasia. BCoV reverse transcription real-time PCR cycle threshold value 16.1. (200X, H&E) (D) BCoV immunohistochemistry demonstrates intense, consistent immunoreactive brown signal in the affected respiratory epithelium (arrow) with intense staining of the slough cell and inflammatory debris of the airway lumen (asterisk). (200X, IHC).

## 4. Discussion

This study assessed the prevalence, temporal trends, demographic risk factors, and co-infections of BCoV detected in diagnostic samples obtained from the respiratory tract of cattle submitted to the ISU VDL for diagnosis of bovine respiratory disease. Overall, 15.38% of samples out of 4505 tested positive for BCoV across different states, production types, and age groups. Previous studies described similar, higher, or lower BCoV detection in North America and worldwide (26, 45–48); however, considering data originated from natural infections and field cases submitted to a diagnostic lab, differences in sampling and testing methods, and study population, precluded accurate comparisons with other BCoV studies.

Age was a significant predictor of BCoV test positivity in this study. The logistic regression model indicated that for each additional day of age, the odds of testing positive for BCoV decreased by approximately 0.15%. This suggests that younger calves were at a higher risk of infection. This finding aligns with previous studies, which have shown that younger calves were more susceptible to respiratory BCoV infections (47). Factors such as variable immune status, overcrowding, transport stress, herd size, presence of co-infections, vaccination status, colder months, and diarrhea within the herd have all been implicated in increasing the risk of BCoV respiratory BCoV infections (26, 49, 50). This age-dependent trend associated with BCoV respiratory infections and clinical disease warrants improving herd health, including appropriate nutrition, reducing stress, vaccination (51), and improving biosecurity measures (52, 53), to reduce the risk of BCoV respiratory infections in young calves. We also explored the temporal trend in BCoV positivity in samples submitted to the ISU-VDL across the study period between January 2020 and November 2025, which revealed a stable pattern in BCoV-positive and negative tests over the study period. The LOESS smoothing line indicated no noticeable fluctuations over time, and the MK test showed a tau value of 0.0102 with a *p*-value of 0.4022, indicating that there was no significant monotonic trend. This finding suggests that there were no major shifts in the prevalence of BCoV over the years of the study, suggesting a stable disease burden in the cattle population. In addition to the demographic and temporal analysis, co-infection patterns among BCoV and other respiratory viral and bacterial pathogens and clustering of strains based on genetic sequences were assessed. Distinct co-occurrence patterns were observed, and viral pathogens were more likely to co-occur with other viral pathogens and bacterial pathogens with other bacterial pathogens. Among the viral pathogens, Bovine Herpesvirus 1 (BHV-1) most frequently co-occurred with Bovine Viral Diarrhea Virus (BVDV), these two viruses with Bovine Respiratory Syncytial Virus (BRSV), and these three viruses with BCoV. Based on the heatmap, the bacterium that was most commonly co-occurring with viral pathogens was *Mannheimia haemolytica*. Among bacterial pathogens, *Histophilus somnus* showed the highest co-occurrence with *Mycoplasma bovis*, followed by *Pasteurella multocida* and *Mannheimia haemolytic*a. This finding suggests complex co-occurrence patterns and potential synergistic effects among viral and bacterial pathogens, which all contribute to the bovine respiratory disease complex (54). Previous studies also described co-occurrence patterns, including BCoV and *H. somni* (55), BCoV and BRSV (46), and several bacterial pathogens and BCoV (56), which were all linked to bovine respiratory diseases.

In addition to co-occurrence analysis, we evaluated clustering of pathogens and quantified their proportions within a cluster. Interestingly, most pathogens (99.55%) clustered into one large cluster that showed a predominance (37-56%) of bacterial pathogens, a medium to low percentage of BCoV and other viral pathogens (1-18%). On the other hand, cluster 2 included samples with a predominance of BCoV (94%) and a high detection rate (56-78%) of viral and bacterial pathogens. These findings suggest that in many cases, bacterial infections are the primary contributors to respiratory disease in the cattle population being studied. However, in a subset of samples, BCoV co-occurred with a high load of viral and bacterial pathogens, potentially contributing to more complex disease presentations with potential severe disease outcomes. This finding is supported by a previous study, where the presence of BCoV was associated with higher mortality in feedlot cattle with respiratory disease (57).

The phylogenetic analysis of complete genome sequences showed that all four whole genomes clustered with other American BCoV strains from Nebraska, Oregon, Kansas, Ohio, and Pennsylvania in group GIIb. The genome organization of all four strains was identical to the BCoV reference strains, except for BCoV-C2, where we reported a small 4.8 kDa non-structural protein (NSP) of 66 nucleotides in length. The BCoV-C6 also demonstrated a small 4.8 kDa NSP of 78 nucleotides in length, like the BCoV respiratory isolate from Nebraska (MARC/2021/01). Furthermore, BCoV-C2 demonstrated a unique small protein (Protein X) of 87 nucleotides between 4.8 kDa and 12.7 kDa that was not observed in the other sequences from C1, C5, and C6. A previous study reported that 4.8 kDa and 4.9 kDa NSP are vestiges of the 11 kDa protein, and even a single nucleotide deletion can give rise to a stop codon (58, 59). Another study reported that truncation of large genomes encoding 4.8 kDa and 4.9 kDa was related to adaptation to human hosts (60), although the truncation was not similar to HCoV-OC43. While another study showed high nucleotide substitution in the 4.8 kDa and 4.9 kDa of BCoV with wild ruminants, which did not support the correlation or natural adaptation (61). The function of these NSPs is not yet completely understood; it is nonessential for viral replication because HCoV-OC43 and porcine hemagglutinating encephalomyelitis virus lack these NSPs (62, 63).

American BCoV and some other strains from China had the mutation N146I and D148G in the spike glycoprotein (3, 64). In this study, we found similar mutations N146I and D148G in the spike glycoprotein of all four BCoV whole genomes (BCoV-C1, C2, C5, and C6). These sites (146 and 148) in the NTD domain near the sialic acid binding region of spike glycoprotein can lead to structure changes (3). Amino acid at position 509, in the receptor-binding domain of the spike glycoprotein. Previous studies reported the mutation N509H (64) and N509T (65), which may lead to BCoV transmission and tissue tropism. Similarly, BCoV-C1, C2, and C6 show the N509T mutations at the spike glycoprotein. These three strains also exhibit another mutation, S501P at the putative receptor-domain (S1-CTD) on the external surface of the spike glycoprotein. However, some of the BCoV from China (HLJ/QQHR-6/2020 and HLJ/QQHR-7/2020) show the S501F mutation at the same position (66). Furthermore, we identified six novel mutations in the BCoV-C5 spike glycoprotein: two in the NTD domain (F232L and H244Y) and two in the CTD domain (A435S and T447I). Similarly, BCoV-C2 has five novel mutations, in which T240M, T617I, and K621E are in the S1 of the spike glycoprotein. Therefore, these mutations could possibly lead to structural changes in the S1 domain and could have a potential effect on receptor binding, viral infectivity, and tissue tropism.

The HE protein has a bimodular structure containing a carbohydrate-binding lectin domain and an active enzymatic esterase domain (67–69). The lectin domain plays a role in viral attachment to the receptors, and the esterase domain performs receptor destruction and virus release. Four amino acids, F211, L212, S213, and N214, present in the lectin domain are important for receptor binding (67). The BCoV-C5 has a mutation at N214Y in the lectin domain of the HE protein. This mutation could affect the lectin activity and affect the viral attachment. A recent study reported the insertion of four amino acids (KATV) at F211 and L212 in the lectin domain, suggesting interspecies transmission (67, 70). In contrast to previously identified mutations, L4P located at the signal peptide region, N49T in the putative esterase domain, and L392I in the membrane-proximal domain of HE protein were also reported in all four isolates (64). The HE protein of all four isolates reported in this study carries a D66G mutation, which has been reported in most BCoV respiratory isolates (71). Another novel S349A mutation was observed in the HE protein of both BCoV-C5 and BCoV-C6, as in the enteric BCoV isolate from Nebraska VDC/2022/06.

In our previous study, we found that BCoV-13 enteric strains from Iowa have a deletion of three amino acids at position 207RAS209 of the nucleocapsid protein (10). Similarly, BCoV NXWZ2310 from China in 2023 (72) shares a similar deletion of 207RAS209 in the nucleocapsid protein. However, BCoV/YNLP/2023 isolates from China have a deletion at 206SRA208 (64), similar to those seen in BCoV from Japan in 2016 to 2017 (30). In this study, we find mutation S209F at the same position in the nucleocapsid protein of the BCoV-C2 strain. Interestingly, BCoV-C2 has another unique mutation, R203C, in the nucleocapsid protein. Furthermore, BCoV-C5 has four unique mutations in the nucleocapsid protein, including Q52L and Q184H, which are located in the RNA-binding domain (NTD). Notably, BCoV-C5 has two unique mutations, S197T and L221V, in the membrane protein. These mutations potentially impact function of the nucleocapsid protein, such as the formation of viral genomic RNA complexes or interactions with membrane proteins.

BCoV is an epitheliotropic virus, and respiratory disease is often associated with varying levels of severity depending on the stage of the disease process, which may be influenced by the time of sample collection in the field (5). Trachea epithelium supports infection and potential replication with BCoV as demonstrated in both experimental and field studies (5, 6), although trachea is not a common sample type submitted for diagnosis of bovine respiratory disease. Prior studies have shown that BCoV is also associated with microscopic lesions and pathology in the lower respiratory tract that is limited to conducting airway epithelium and some degree of epithelial attenuation, necrosis, or proliferation, as demonstrated in the examples provided in this study (Figure 12). In addition, the magnitude of lung pathology may be affected by the stage of the disease process and level of replication in the lung, as suggested by semi-quantitative cycle threshold (Ct) values reported from RT-PCR. The histopathology of the conducting airways presented in Figure 12A and B demonstrates subacute, regenerating lesions and limited IHC signals associated with a Ct 29.1, in contrast to Figure 12C and D, where epithelial attenuation and acute lung lesions were associated with a Ct 16.1 and intense IHC signals in the epithelium. Magnitude of viral replication and the stage of disease process should be interpreted with caution when based on Ct values and using samples collected from the field where BCoV strain, time, and level of challenge, presence of co-infections, and environment are uncontrolled but may affect diagnostic outcomes or results. Regardless, our results indicate the detection of BCoV by RT-PCR and IHC when associated with histopathological lung lesions, even in the presence of co-infections, support a diagnosis of BCoV respiratory disease despite its frequent detection in enteric cases (4, 73) or even when calves are unaffected by respiratory disease (49, 74).

## Acknowledgement

We thank the Iowa State University Veterinary Diagnostic Laboratory (ISU-VDL) for providing samples and for their technical assistance with the histopathology and immunohistochemistry testing.

## Data availability statement

Data is available from the authors upon reasonable request.

## Author contribution statement

AUS; Software, data analysis, writing the first draft, writing the final manuscript. **CV;** epidemiological data analysis, statistical analysis, writing the first draft, writing the final manuscript. **PG;** conceptualization, sample collection, histopathology, IHC, writing first draft, writing the final manuscript. **MGH;** conceptualization, monitoring, funding acquisition, writing the first draft, writing the final manuscript.

